## Supplementary figures and images for "Genomic fingerprints of the world’s soil ecosystems"

### FigS1

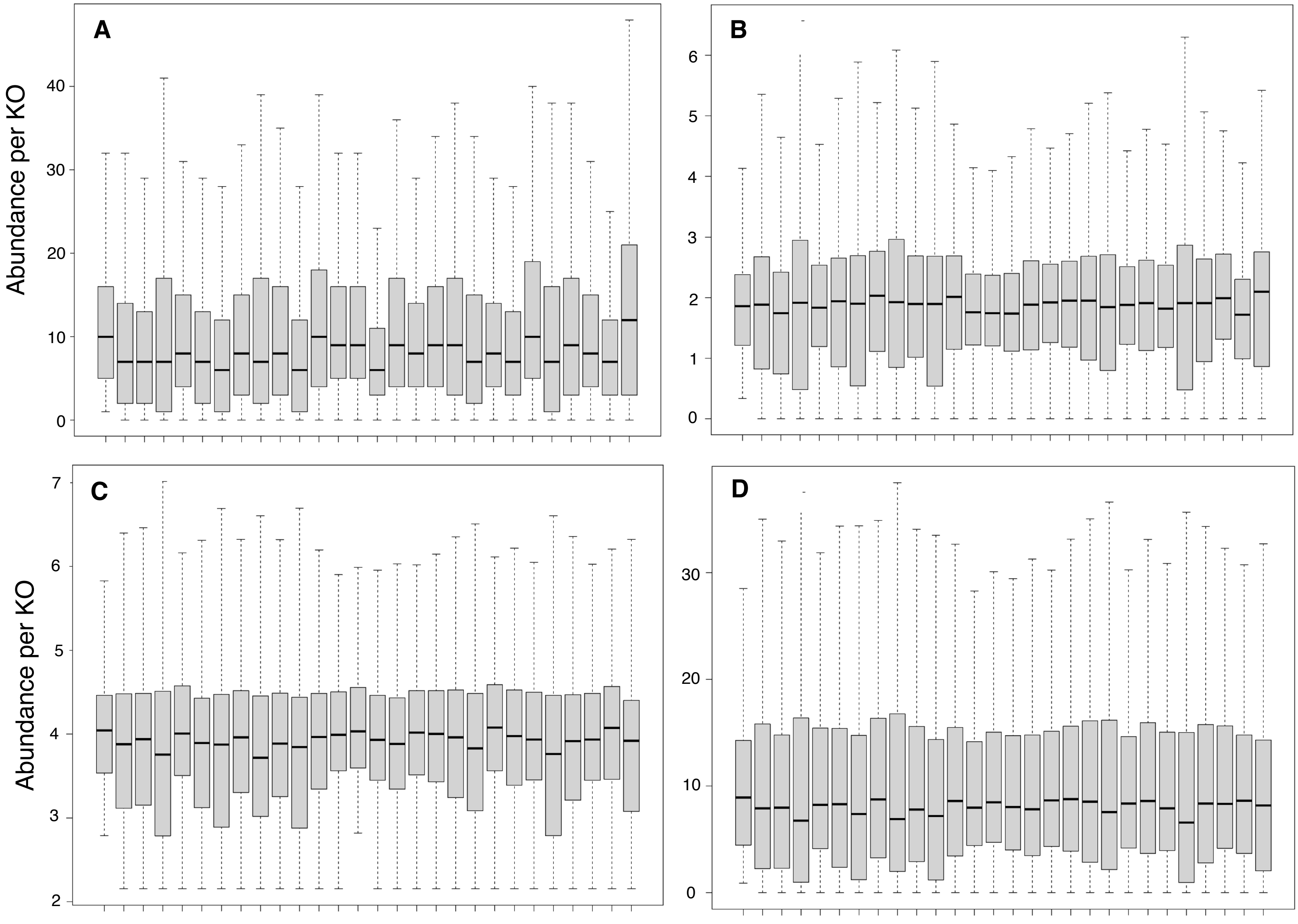

### FigS2

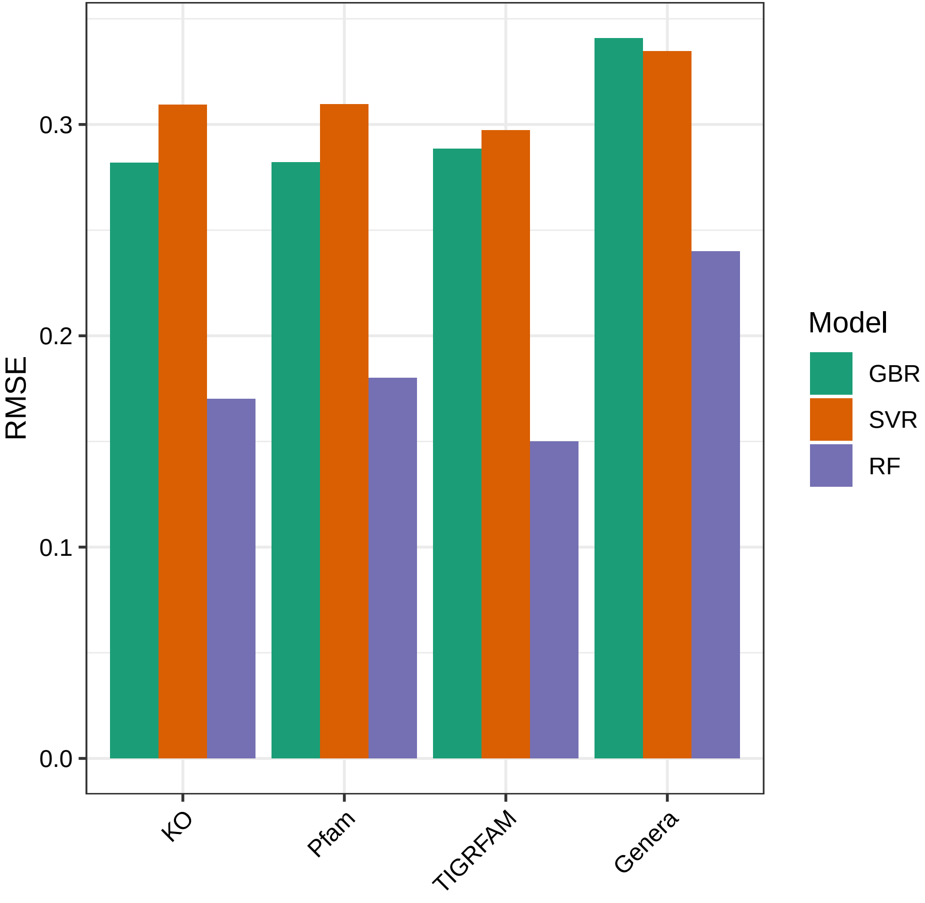
