## Supplementary material for "Genomic fingerprints of the world’s soil ecosystems": Legends for Fig S1-2

**Figure S1. Abundances across different normalization procedures.** (A) Raw data, (B) CSS, (C) DESeq2, and (D) TMM. KO data from selected samples are shown as an example. Each box represents one sample. Sample names are removed for visual clarity. Upper and lower hinges of the box plots represent the 75th and 25th percentiles and whiskers represent 1.5 times the 75th and 25th percentiles, respectively. Colors coincide with labels on the x-axis.

**Figure S2. Comparison of machine learning algorithms.** Model performance for parsing KO across bulk density is shown. Results for all other comparisons followed the same pattern (i.e., random forest models consistently had the lowest root mean square error). All models were constructed using a 50%-50% split between testing and training datasets with 10-fold cross validation. The root mean square error (RMSE) of gradient boosted regression (GBR), support vector regression (SVR), and random forest (RF) algorithms is shown in green, red, and purple, respectively.
